# A codon model of nucleotide substitution with selection on synonymous codon usage

**DOI:** 10.1101/007849

**Authors:** Laura Kubatko, Premal Shah, Radu Herbei, Michael A. Gilchrist

## Abstract

The quality of phylogenetic inference made from protein-coding genes depends, in part, on the realism with which the codon substitution process is modeled. Here we propose a new mechanistic model that combines the standard M0 substitution model of Yang (1997) with a simplified model from Gilchrist (2007) that includes selection on synonymous substitutions as a function of codon-specific nonsense error rates. We tested the newly proposed model by applying it to 104 protein-coding genes in brewer’s yeast, and compared the fit of the new model to the standard M0 model and to the mutation-selection model of Yang and Nielsen (2008) using the AIC. Our new model provided significantly better fit in approximately 85% of the cases considered for the basic M0 model and in approximately 25% of the cases for the M0 model with estimated codon frequencies, but only in a few cases when the mutation-selection model was considered. However, our model includes a parameter that can be interpreted as a measure of the rate of protein production, and the estimates of this parameter were highly correlated with an independent measure of protein production for the yeast genes considered here. Finally, we found that in some cases the new model led to the preference of a different phylogeny for a subset of the genes considered, indicating that substitution model choice may have an impact on the estimated phylogeny.

## Introduction

Successful phylogenetic inference based on protein-coding genes relies on use of an appropriate model of sequence evolution. Models in current use are of one of two classes: models that use information in the nucleotides only, without taking into account information about codons or amino acids (e.g., the GTR model (Tavare, 1986; Yang, 1994; Zharkikh, 1994) and its submodels), and codon-based models that are designed to incorporate information about rates of change between pairs of amino acids (e.g., Yang et al. (1998); Kosiol et al. (2007)). Codon-based models can be further classified into empirical and mechanistic models (Kosiol et al., 2007). Empirical models use the observed frequencies of changes in state observed within large data sets to specify the rate of change used in the model. These estimated rates are assumed to be applicable to a broad set of sequence data sets, and thus parameters are not generally estimated separately for a particular data set. Empirical models have been widely used for modeling the amino acid substitution processes (Dayhoff and Eck, 1968; Dayhoff et al., 1972, 1978; Jones et al., 1992; Whelan and Goldman, 2001; Jones et al., 1994; Goldman et al., 1996, 1998; Adachi and Hasegawa, 1996; Adachi et al., 2000; Dimmic et al., 2002; Yang, 1994).

Mechanistic models, on the other hand, specify an explicit model for the evolutionary process using features such as selective pressures acting on certain types of changes and the varying frequencies of codons in the data. Nearly all codon models in common use are mechanistic (but see Kosiol et al. (2007) and references therein). In general, codon substitution models are based on Markov models of the rates of nucleotide substitution and typically include a parameter to quantify differences in the rates of synonymous versus nonsynonymous substitutions. The magnitude of this parameter, << 1 or >> 1, is often taken as evidence for protein-level stabilizing or diversifying selection, respectively (Goldman and Yang, 1994; Yang and Nielsen, 1998; Yang and Bielawski, 2000; Yang et al., 2000). Extensions of this approach have been proposed to test for selection at specific sites in the sequence, in specific lineages of the phylogeny, or both (Yang et al., 2000; Wong et al., 2004; Massingham and Goldman, 2005; Yang and Nielsen, 1998, 2002).

Within the last ten years, these basic models have been extended in several important directions to attempt to capture the complexities in the codon substitution process in the presence of selection. For example, Kosakovsky Pond and Muse (2005) allowed the rates of synonymous and nonsynonymous substitutions to vary across position in the sequence, while Mayrose et al. (2007) constructed a family of models to allow both variation in synonymous and nonsynonymous rates across sites and site-to-site dependence in rates via a first-order Markov process. Nielsen et al. (2007) and Zhou et al. (2010) used the idea of “preferred” and “non-preferred” synonymous substitutions, where a synonymous substitution is either preferred or non-preferred at a site depending on selective forces specific to that location. A primary emphasis in both of these studies is the development of ways to measure selection along the genome and across branches of a phylogeny. Several recent models have also separated the substitution rate into components due to the mutational process and the selection process. For example, Yang and Nielsen (2008) introduced the FMutSel model, in which each codon is assigned a fitness parameter; differences in the fitness parameters between two codons are used to specify the substitution rates in the Markov matrix, by modifying the rates specified by the standard mutation models. Similarly, Rodrigue et al. (2010) developed a model in which amino acid propensity scores are used to estimate scaled selection coefficients that are then used to specify substitution rates.

In this paper, we propose a new mechanistic codon-based substitution model that takes into account selection on codon usage of a gene. Our model is based on the M0 model in PAML (Yang, 1997) but where the substitution rate between codons, including synonymous changes, is modified by the substitution’s effects on protein production costs and the average protein production rate of the gene. We calculate effects of a codon substitution on the production cost of a protein using a model of protein translation based on the movement of the ribosome along the mRNA of a gene (Gilchrist and Wagner, 2006; Gilchrist, 2007). The production cost includes the effects of nonsense (a.k.a. processivity) errors which result in premature termination of the translation of a protein. More specifically, changes in protein production costs are due to presumed inter-codon variation in nonsense error rates.

We fit our model to empirical data for 104 genes from 8 yeast species, and compare the fit our model to several of the codon models in common use, such as the M0 model in PAML and the FMutSel model of Yang and Nielsen (2008). We also examine the effect of the substitution model on phylogenetic inference by considering which of two competing phylogenies for these 8 yeast species is preferred by various models.

## New Approaches

### Background: Codon Substitution Models

To motivate development of our method, we review the codon substitution models in common use. Specifically, we give the details of the M0 model implemented in the program PAML. This model uses a continuous-time Markov model for the substitution process between codons in a protein-coding gene. The states in the Markov process are the 61 sense codons (stop codons are not included). The model is then specified by a 61 × 61 matrix **Q**, whose entries *Q_ij_* give the instantaneous rate of substitution of codon *i* with codon *j* and satisfy the constraint *Q_ii_* = Σ*_j≠i_ Q_ij_* . The probabilities of substitution of codon *i* with codon *j* over time *t* can then be found by solving the matrix differential equation **P**′(*t*) = **QP**(*t*) with initial condition **P**(0) = **I**, which yields **P**(*t*) = exp*{***Q***t}*. Thus specification of **Q** and the stationary frequencies of the codons is sufficient to compute the substitution probabilities that will be used in modeling the codon mutation process.

Define codon *i* to be *i*_1_*i*_2_*i*_3_ and codon *j* to be *j*_1_*j*_2_*j*_3_, where *i_k_, j_k_ ∈ {A, C, G, T }* for *k* = 1, 2, 3. The M0 model specifies **Q** as follows:

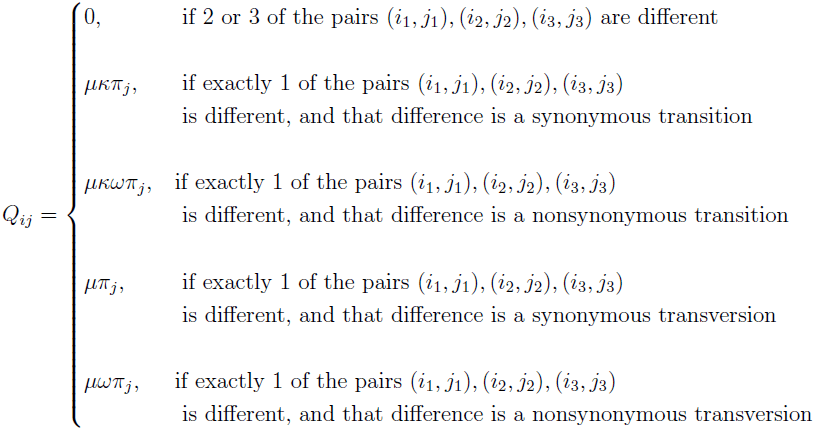

In the above expression, the parameter *κ* allows for a different rate of substitution for transitions versus transversions. The *ω* parameter is used to specify a different rate for synonymous and nonsynonymous substitutions. These parameters are typically estimated from the data, and are often used to provide insight into the mode of evolution of a particular gene. For example, an *ω >* 1 indicates positive selection, while an *ω <* 1 indicates negative selection. When *ω* is not significantly different than 1, there is no evidence that the gene is under selection. The *π_j_* parameters give the frequency of each of the 61 possible codons at equilibrium. There are several options for setting these parameters: they may all be set to be equal (the ‘Fequal’ model in PAML), they may be estimated as 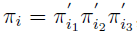, where 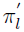 refers to the empirical frequency of nucleotide *l* for that gene (the ‘F1 × 4’ model in PAML), they may be estimated as 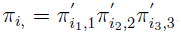, where 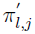 refers to the empirical frequency of nucleotide *l* at codon position *j* (*j* = 1, 2, 3) for that gene (the ‘F3 × 4’ model in PAML), or empirical estimates of each of the 61 codons may be used (the ‘Fcodon’ model in PAML). Finally, the parameter *µ* is set so that 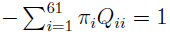, which scales time along the tree to be in units of expected numbers of nucleotide substitutions per codon. Finally, we note that the model above satisfies the condition of time reversibility, i.e., *π_i_Q_ij_* = *π_j_ Q_ji_* for all *i* and *j*. Given a particular **Q** matrix, **P**(*t*) can be calculated using standard numerical algorithms (see (Yang, 2006; Moler and Van Loan, 2003; Golub and van Loan, 1996) for details).

### Background: Modeling Protein Translation

Using the approach developed in Gilchrist (2007), we can calculate the cost of producing a complete and functional protein product in the face of nonsense errors. Briefly, the production cost of a protein is equivalent to the ratio of the expected cost to the expected benefit, i.e. functionality, of a protein produced from a given coding sequence (Gilchrist et al., 2009). Functionality is defined on a 0 to 1 scale relative to the functionality of a complete and error-free protein produced from the coding sequence. This allows us to avoid having to consider the specific biological role a protein plays.

Let *q_j_* represent the probability of elongation of a codon of type *j*, where *j* ranges over the 61 sense codons (*AAA, . . ., TTT*). Let the codon at position *i* of a gene of length *n* be represented by *c_i_*, where *i* = 1, 2*, . . ., n*. The probability that the codon at the *i*^th^ position, will be successfully translated is given by *p_i_* = *q_ci_*, and thus 1 *-p_i_* represents the probability that a nonsense error occurs at that codon. It follows that the expected cost of translating a coding sequence 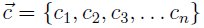 of length *n*, is

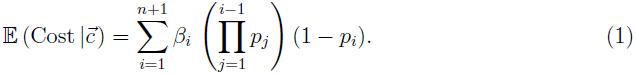

where *β_i_* represents the energetic investment, in Adenosine Tri-Phosphate molecules (ATPs), of translating *i* codons successfully. Note that the summation is taken up to *n* + 1 to account for the stop codon and (1 *-p_n_*_+1_) = 1 by definition. In general, *β_i_* = *a*_1_ + *a*_2_*i* where *a*_1_ and *a*_2_ represents the cost of translation initiation and protein elongation, respectively, and *a*_1_ = *a*_2_ = 4ATP.

In order to calculate the expected functional benefit of a protein produced from a coding sequence 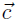 We begin with the simplifying assumption that truncated proteins have zero functionality. Because we measure functionality on a relative scale, such that a complete, error free protein has a functionality of 1, it follows that this expected functionality of a gene is simply equal to the probability of producing a complete protein such that,

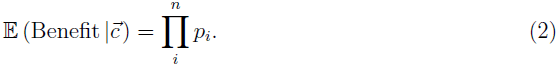

Then the expected cost-benefit of protein production for a given codon sequence, which we call the protein production cost for brevity, is 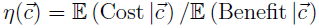. By including the denominator into the summation term, we can re-write this as

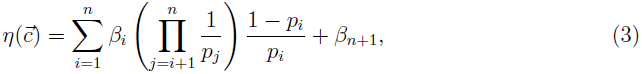

(see Gilchrist et al. (2007) for more details).

Due to the non-linear nature of the protein production cost *η*, the effect of changing codon *c_k_* to codon 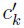 at position *k* on *η* will depend on the codons at the other sites. In other words, the effect of a codon substitution on *η* is not independent between sites. To take such inter-dependence explicitly into account when trying to formulate the substitution rate matrix **Q** would be unfeasible computationally. However, given that the probability of a nonsense error at a codon, 1*-p_i_*, is much smaller than the elongation probability *p_i_*(Gilchrist, 2007), the first-order approximation of Equation 3 can be written as

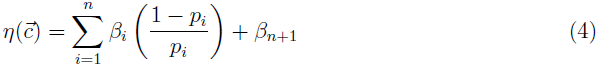

Since different codons have different elongation probabilities, 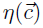 will vary between alleles of a coding sequence. More specifically, if two alleles, 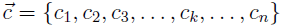 and 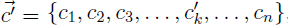, differ at codon *k* and 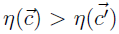 then allele 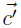 should be *favored* by natural selection because of its lower protein production cost. Based on Equation 4, the cost of substituting codon *c_k_* with 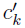 depends only on the position *k* and the elongation probabilities of the two codons involved, *p_k_* and 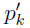:

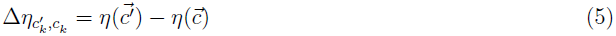

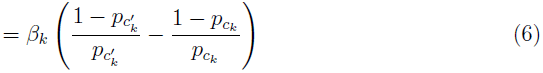

The goal of our work is to incorporate these differences in protein production costs due to inter-codon variation in elongation probabilities into our substitution model.

### A New Codon Substitution Model: MutNSE

The main idea behind our new model, which we call MutNSE to indicate that the model incorporates both the mutation process and nonsense errors, is to combine the features of the models described in the previous two subsections in order to incorporate two separate features of the process of the codon substitution in an explicit manner. These features are (1) the typically modeled rate biases (transition/transversion and synonymous/nonsynonymous), and (2) the change in the probability of the protein being ultimately produced following substitution of one amino acid by another. For (2), we note that the probability of successful protein production following codon substitution varies throughout the sequence. We take this aspect of the model into account by specifying a different instantaneous rate matrix **Q** for each codon position *k* in the sequence as follows:

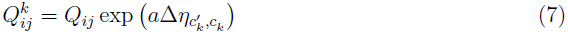

where *Q_ij_* refers to the substitution rates in the standard codon substitution model of choice and 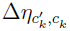 is defined in (6). Selection on codon usage has been known to vary between genes. In particular, codon usage in genes with low expression is driven by patterns of mutation biases, while codon usage in high expression genes is primarily driven by natural selection (Sharp and Li (1986), Bulmer (1991), Shah and Gilchrist (2011), Wallace et al. (2013)). The parameter *a* scales the contribution of selection on codon usage to the overall substitution rate and is a free parameter in our model that we estimate. When there is no selection on codon usage, then *a ≈* 0 and the model reduces to the standard codon substitution model. When *a* is large, the substitution probabilities depend primarily on changes in protein production costs *η*.

Our model makes certain simplifying assumptions about the evolution of tRNA copy number and expression levels of genes. Similar to other codon-based models of protein evolution that incorporate selection on individual codons, we assume that the selection on synonymous codons remains fairly constant across the phylogenetic breadth of organisms under consideration. In our mechanistic framework, this translates to the assumption that the variation in tRNA copy numbers and expression levels of genes is quite small across the phylogeny.

Our model requires calculation of separate **Q** and **P**(*t*) matrices for each codon position *k* in the sequence. Because this is computationally intensive, we have developed a Graphical Processing Unit (GPU) implementation of this step of the method. In the past few years, GPUs have made a significant contribution to scientific computing due to their ability to perform massive parallel calculations. GPU-based algorithms have now made their way into mainstream computing in many fields, see for example, Suchard and Rambaut (2009), Suchard et al. (2010), Lee et al. (2010), Cron and West (2011), Herbei and Kubatko (2013). The work described in this paper has been implemented and tested initially on a NVIDIA Tesla C2075 GPU while the bulk of the computing was done on the OAKLEY cluster at the Ohio Supercomputing Center (www.osc.edu), which has 128 Tesla M2070 GPUs.

For this work we require fast, parallel evaluation of the transition matrix **P**(*t*) = exp(*t***Q**). Due to the complexity of our approach and the size of the problem (a typical gene has *∼*300-400 codons), a sequential evaluation/approximation for each of the required matrix exponentials is far too inefficient. The speed gained by distributing this computing aspect to the GPU comes from the ability of the user to pre-specify a computing array of independent threads. Each thread will evaluate a matrix exponential for a given combination of site/branch length/rate matrix. It is well known that exact evaluation of a matrix exponential can only be done in a few particular cases, while in general, an approximation algorithm is required. A suite of algorithms and their performance is discussed in Moler and Van Loan (2003); Golub and van Loan (1996). For this work, we implemented Method 3 described in Moler and Van Loan (2003), see also Algorithm 11.3-1 of Golub and van Loan (1996). Note that, for a matrix *A*,

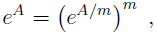

for any integer *m* ≥ 1. When *m* is selected to be a power of 2, the exponential *e^A^* is then obtained through repeated squaring. For the matrix *e^A/m^*, we use the Padé approximation, see Moler and Van Loan (2003), page 9. This approach is characterized as “the only generally competitive series method” (Moler and Van Loan, 2003) and requires basic matrix algebra (matrix scaling/multiplication), thus it is very suitable for parallel computing. The GPU implementation results in an approximately 40-fold reduction in computation time over a standard CPU application.

### Model Comparison

We compare the fit of our proposed model with several models in PAML using the AIC (Burnham and Anderson, 2002). Denoting the maximized likelihood under our new model by *L̂_M utN SE_* and the maximized likelihood under the model in PAML by *L̂_PAML_*, the AIC for each model is computed using

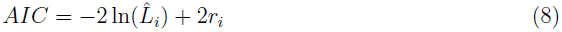

where *i* refers to either the MutNSE model or one of the models in PAML, and *r_i_* is the number of parameters in model *i*.

The models we consider are listed in Table 1. In particular, we consider the MutNSE model with four different choices for the codon frequencies (‘Fequal’, ‘F1x4’, ‘F3x4’, and ‘Fcodon’), as well as the M0 model in PAML with the same four choices for the codon frequencies. Finally, we consider the mutation-selection model of Yang and Nielsen (2008) as implemented in PAML (FMutSel). We compare the MutNSE model against the corresponding M0 model, as well as against the FMutSel model, resulting in seven separate comparisons (Table 2).

**Table 1:** Models considered in this study, along with number of parameters estimated. The notation *π_i_* refers to nucleotide frequencies, and involves 3 parameters since 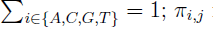 refers to nucleotide frequencies at each codon position *j* = 1, 2, 3, and involves 9 free parameters; *π_J_* refers to codon frequencies, and involves 60 free parameters, corresponding to the 61 sense codons. In the number of parameters to be estimated listed in the table below, we do not include branch length parameters, since this number will be the same under all models.

| Model | Parameters estimated | Total number of parameters |
| --- | --- | --- |
| M0, equal frequencies | $\kappa, \omega$ | 2 |
| MutNSE, equal frequencies | $a, \kappa, \omega$ | 3 |
| M0, F1x4 | $\pi_i, \kappa, \omega$ | 5 |
| MutNSE, F1x4 | $\pi_i, a, \kappa, \omega$ | 6 |
| M0, F3x4 | $\pi_{i,j}, \kappa, \omega$ | 11 |
| MutNSE, F3x4 | $\pi_{i,j}, a, \kappa, \omega$ | 12 |
| M0, Fcodon | $\pi_J, \kappa, \omega$ | 62 |
| MutNSE, Fcodon | $\pi_J, a, \kappa, \omega$ | 63 |
| FMutSel | $\pi_i, \pi_J, \kappa, \omega$ | 65 |

**Table 2:** Number of yeast genes (of 104 total genes) for which the newly proposed model (MutNSE) was preferred over an existing model. The second two columns list the models being compared (see Table 1) and the fourth column gives the number of yeast genes for which the MutNSE model was preferred using AIC.

| Comparison | Model 1 | Model 2 | Number of Times Model 1 Was Preferred |
| --- | --- | --- | --- |
| 1 | MutNSE, equal frequencies | M0, equal frequencies | 93 |
| 2 | MutNSE, F1x4 | M0, F1x4 | 91 |
| 3 | MutNSE, F3x4 | M0, F3x4 | 88 |
| 4 | MutNSE, F3x4 | M0, Fcodon | 26 |
| 5 | MutNSE, Fcodon | M0, Fcodon | 27 |
| 6 | MutNSE, Fcodon | FMutSel | 0 |
| 7 | MutNSE, F3x4 | FMutSel | 2 |

### Application to Yeast Protein-Coding Genes

We applied our model to the 104-gene data set of Rokas et al. (2003). We fixed the tree topology to be the ML tree found by these authors (see Figure 1(a)), and then estimated the MLEs of all model parameters (e.g., *κ, ω, a* and the branch lengths) along this fixed tree. We compared the fit under various models using the AIC, as described above. We also compared the parameter estimates for *κ* and *ω* under the new model and under the M0 model.

**Figure 1:**
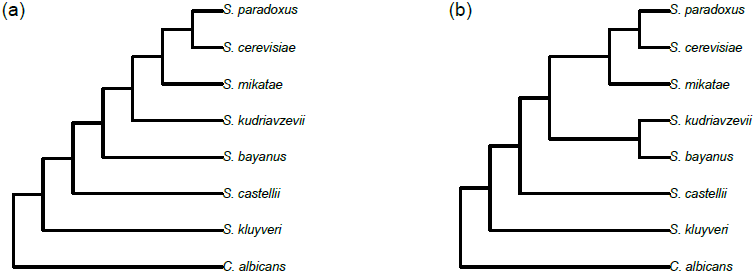
Two phylogenetic trees for the yeast data. The tree in (a) was found by Rokas et al. (2003) to be the ML tree for the concatenated data. The tree in (b) has been proposed by several authors (see, e.g., Edwards et al. (2007)) to be a plausible species-level phylogeny for yeast.

Because the parameter *a* in the MutNSE model can be interpreted as a measure of the extent to which selection on codon usage contributes to the overall substitution rate, we hypothesized that *a* should be correlated with the rate of protein production. To examine this, we obtained protein production rates, *ϕ*, for all *S. cerevisiae* genes from Yassour et al. (2009). These values indicate the average rate of protein production and were estimated using mRNA abundances (MacKay et al., 2004) and ribosome occupancy on mRNAs (Arava et al., 2003). We assume strong purifying selection on protein production rate of the genes considered here. Thus *ϕ* values estimated in yeast are used as proxies for *ϕ* across the phylogeny.

Finally, we wanted to examine whether use of the new model would impact the preferred topology under the maximum likelihood criterion. To examine this, we considered two candidate topologies (Figure 1). Various studies using these data in different modeling frameworks have preferred one or the other of these trees (see, for example, Edwards et al. (2007)). For each gene, we obtained the maximized value of the likelihood under both the MutNSE model and the models in PAML. We counted the number of genes for which a different tree was preferred based on the likelihood under the various models.

## Results

The model comparison results for each of the seven comparisons are shown in Table 2. The MutNSE model is preferred at least 85% of the time when it is compared to the corresponding M0 model with a relatively simple model for estimating the codon frequencies (i.e., ‘Fequal’, ‘F1x4’, and ‘F3x4’). However, when codon frequencies are estimated using the empirical frequencies, the MutNSE model is only preferred about 25% of the time. When the mutation-selection model FMutSel is used, the MutNSE model no longer provides better fit, except for a couple of genes.

Figure 2 shows various relationships between parameter estimates for both the MutNSE and the M0 models in the case when the ‘F3x4’ estimates of codon frequencies were used (results for the other cases are similar and are not shown here). Figures 2(a) and 2(b) compare the estimates of *κ* and *ω* under the two models, with the line indicating equality of the estimates. In both cases, the estimates under the MutNSE model and under the M0 model are highly correlated, with a slight bias toward higher estimates of both parameters under the M0 model. In particular, there are several genes for which the estimates of *ω* are larger under the M0 model. This may be an important finding, as this parameter is often used as a test for and measure of the strength of either purifying or diversifying selection across a gene.

**Figure 2:**
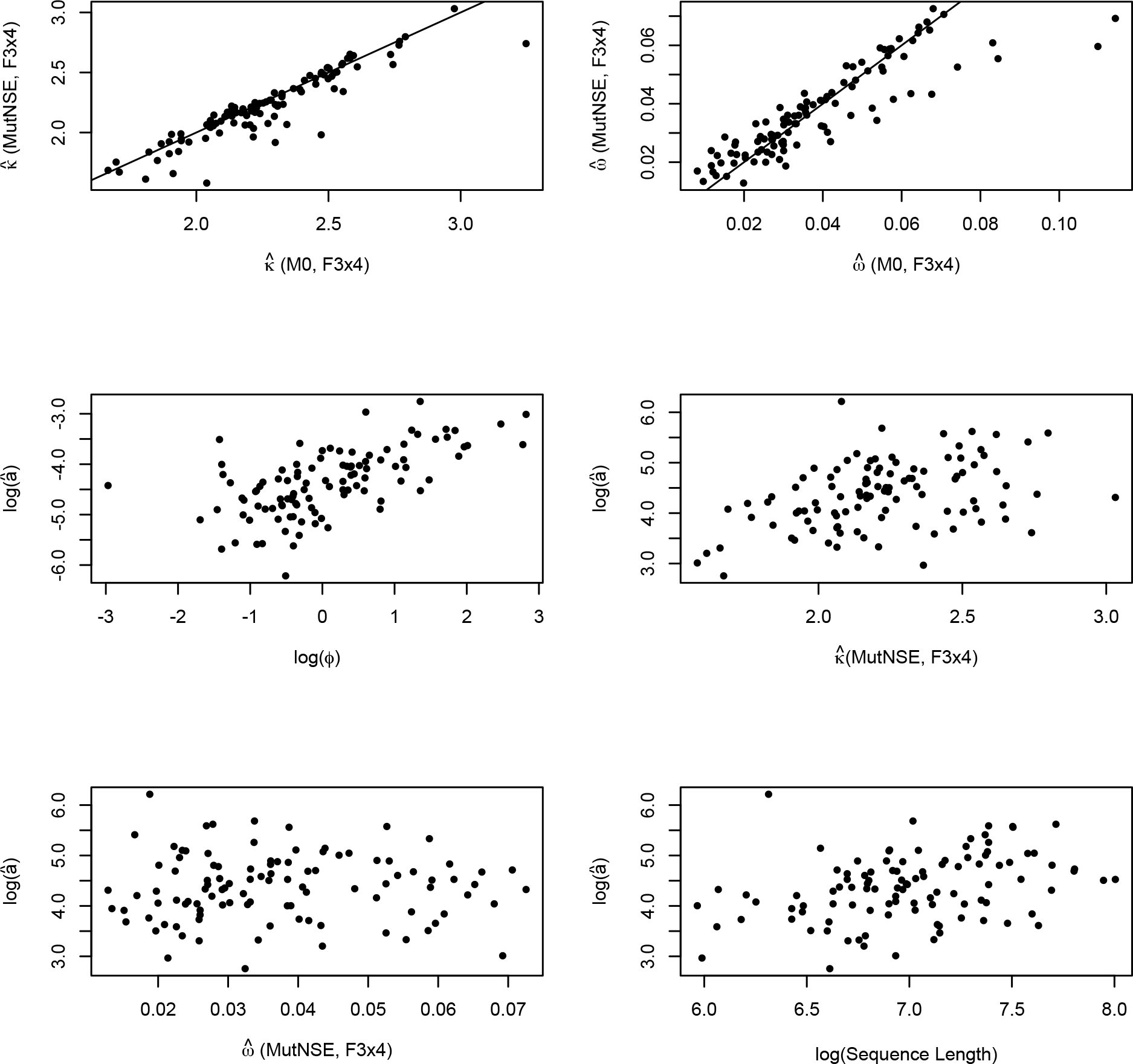
Parameter estimation results for the yeast genes for the MutNSE, F3x4 model and the M0, F3x4 model. (a) Comparison of the values of the parameter *κ* estimated for the MutNSE and for the M0 model. (b) Comparison of the values of the parameter *ω* estimated for the MutNSE model and for the M0 model. In both (a) and (b), the line has slope 1 and represents equality of parameter estimates in the two models. (c) Plot of the estimated value of *a*, which determines the relative importance of codon usage in driving sequence evolution, versus an independent estimate of the rate of protein production, *ϕ* (see text for details on how the estimates of *ϕ* were obtained). (d) Plot of the estimated values of *a* versus the estimated value of *κ* in the MutNSE model. (e) Plot of the estimated values of *a* versus the estimated value of *ω* in the MutNSE model. (f) Plot of the estimated values of *a* versus sequence length in base pairs (bp).

Figure 2(c) shows that the estimated value of the MutNSE model parameter *a* is highly correlated with observed protein production rate (*ϕ*) of yeast genes. This suggests that selection against nonsense errors plays a significant role in affecting the evolutionary rate of highly expressed genes.

Figure 2(d), (e), and (f) examine the relationship between the estimated values of *a* and of other characteristics of the model, specifically the estimated values of *κ*, of *ω*, and the sequence length, respectively. We expect no relationship among these quantities and that is, in fact, what is observed in these plots.

To examine consistency of model preference across genes, we compared the model selected across all genes. Figure 3 shows genes (x-axis) for which the MutNSE model is preferred (indicated with a colored dot) across the various model comparisons (y-axis; height of the points corresponds to the comparison number in Table 2). It is clear that there is consistency across comparisons overall, though there are also differences. In particular, results for the first three comparisons (in which the MutNSE model is preferred the majority of the time) are very consistent, while there is less consistency across comparisons 4 and 5 (in which the MutNSE model is only preferred 25% of the time).

**Figure 3:**
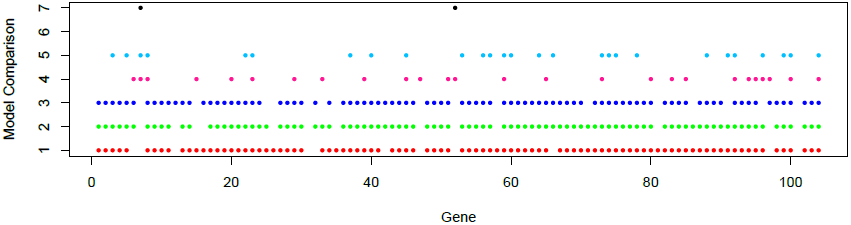
Comparison of model selection results across genes (x-axis) and across comparisons (y-axis; see Table 2 for comparison numbers). For all comparisons, a colored dot indicates preference for the MutNSE model over the relevant existing model.

We also compared the MutNSE model to the M0 model with ‘F3x4’ in terms of which of the two topologies in Figure 1 was preferred under each model using the likelihood value. The MutNSE model had a higher likelihood for the tree in Figure 1 (a) for 42 of the 104 genes, while the M0 had a higher likelihood for only one of the 104 genes. This means that for 41 of the 104 genes, the likelihood criterion would order the two trees in Figure 1 differently under the MutNSE model versus the M0 model. Thus, model choice can impact estimation of the phylogeny.

Finally, we note that because the MutNSE model is site-specific, likelihood computations using the model are non-trivial. The GPU computing machinery used here is crucial to obtaining phylogenetic estimates in reasonable time. Using the OAKLEY Cluster at the Ohio Supercomputer Center, each likelihood evaluation takes under a minute, and full optimization of all model parameters (including branch lengths) along a fixed tree requires between 5 minutes and 2 hours for most genes. We point out that the only step of our implementation that takes advantage of GPU computing is the computation of transition probabilities for each site. Likelihood computation across a tree has also been implemented in a GPU framework (Ayres et al., 2012) and this would speed computations even further.

## Discussion

Overall, our results indicate that incorporating selection on synonymous codon usage is an important component of a codon substitution model, as has been noted by others (Yang and Nielsen, 2008; Nielsen et al., 2007; Zhou et al., 2010; Rodrigue et al., 2010). We found that when simple models of codon frequencies were used, our MutNSE model was preferred over the M0 model for the majority of data sets (*>* 85% of genes in the yeast data set). However, when empirical codon frequencies were used in both models, the new model was preferred for only about 25% of the genes, and when a model that incorporates both mutation and selection was used (the FMutSel model), our model was generally not preferred using the AIC. This is not completely unexpected, because the FMutSel model is parameterized so that estimates of the selection parameters are obtained empirically, while our MutNSE model incorporates the effect of codon usage via the inclusion of elongation probabilities that are obtained independently and are fixed across genes. However, the set of comparisons made here highlights the importance of realistic models for both codon frequencies and for the process of selection. This is particularly apparent by noting that the MutNSE model preferred the tree in Figure 1(b) over that in Figure 1(a) for 42 of the 104 genes, while the corresponding M0 model only preferred the tree in Figure 1(b) for a single gene. Thus the choice of substitution model can have an important impact on phylogenetic inference.

An important feature of our MutNSE model is that it is able to accurately predict the level of protein production. For genes with high expression (*φ*), we find that selection on codon usage against translation errors is a significant determinant of evolutionary rate (see also Drummond et al. (2006)). This observation is particularly important given that genes used in building phylogenies tend to have a broad phylogenetic breadth, and are highly expressed (Nei et al., 1997, 2000; Eiŕın-Ĺopez et al., 2004). Thus, it is essential to develop models of codon substitution that explicitly take into account the effects of selection on synonymous codons and how they change with gene expression. We expect that such models should improve both the reliability and accuracy of the parameters estimated as part of phylogenetic analyses, especially in terms of evaluating whether the ratio of synonymous to non-synonymous substitutions is consistent with stabilizing vs. diversifying selection on the amino acid sequence of a gene.

The model presented here takes into account selection on synonymous codon usage against premature termination. However, patterns of codon usage are also under selection pressures for translation accuracy (Akashi, 1995; Drummond and Wilke, 2008, 2009) and efficiency (Bulmer, 1991; Plotkin and Kudla, 2011; Shah and Gilchrist, 2011). Although the relative importance of these pressures are actively debated in the field, the selective advantage of a synonymous codon for both efficiency and accuracy has been shown to be correlated with its tRNA abundance (Shah and Gilchrist, 2011; Wallace et al., 2013), as has been assumed here. Because we incorporate selection on codon usage in a mechanistic manner, expanding our model to include these additional selective forces is possible in future implementations. Such extensions should not only improve our ability to reconstruct evolutionary relationships based on DNA sequence data, but also potentially extract additional information on key parameters related to the protein translation process itself such as codon-specific nonsense error rates or ribosome pausing times.

## Acknowledgements

We thank Ziheng Yang for providing information concerning the parameterization of the models in PAML. This work was supported by the National Science Foundation (DEB0918195 to L.S.K.; MCB1120370 to M.A.G.; and EF0832858 to the National Institute for Mathematical and Biological Synthesis (NIMBioS), which supported M.A.G. and P.S.); the University of Tennessee Knoxville, NIMBioS, and a Tennessee Science Alliance Joint Directed Research and Development grant (to M.A.G.); and the Alfred P. Sloan Foundation, Burroughs Wellcome Fund Career Award, David & Lucille Packard Foundation Fellowship (to J. B. Plotkin and used to support P.S.). We also acknowledge support from the Ohio Supercomputer Center and the Mathematical Biosciences Institute (MBI) at The Ohio State University.

